# Exposure route, sex, and age influence disease outcome in a golden Syrian hamster model of SARS-CoV-2 infection

**DOI:** 10.1101/2021.06.12.448196

**Authors:** Bryan D. Griffin, Bryce M. Warner, Mable Chan, Emelissa J. Mendoza, Nikesh Tailor, Logan Banadyga, Anders Leung, Shihua He, Amrit S. Boese, Jonathan Audet, Wenguang Cao, Estella Moffat, Lauren Garnett, Kevin Tierney, Kaylie N. Tran, Alixandra Albietz, Kathy Manguiat, Geoff Soule, Alexander Bello, Robert Vendramelli, Jessica Lin, Yvon Deschambault, Wenjun Zhu, David Safronetz, Heidi Wood, Samira Mubareka, James E. Strong, Carissa Embury-Hyatt, Darwyn Kobasa

**Author notes:** These authors contributed equally (Bryan D. Griffin, Bryce M. Warner).

## Abstract

The emergence of Severe acute respiratory syndrome coronavirus 2 (SARS-CoV-2) and the resultant pandemic of coronavirus disease 2019 (COVID-19) has led to over one hundred million confirmed infections, greater than three million deaths, and severe economic and social disruption. Animal models of SARS-CoV-2 are critical tools for the pre-clinical evaluation of antivirals, vaccines, and candidate therapeutics currently under urgent development to curb COVID-19-associated morbidity and mortality. The golden (Syrian) hamster model of SARS-CoV-2 infection recapitulates key characteristics of severe COVID-19, including high-titer viral replication in the upper and lower respiratory tract and the development of pathogenic lesions in the lungs. In this work we examined the influence of the route of exposure, sex, and age on SARS-CoV-2 pathogenesis in golden hamsters. We report that delivery of SARS-CoV-2 primarily to the nasal passages (low-volume intranasal), the upper and lower respiratory tract (high-volume intranasal), or the digestive tract (intragastric) results in comparable viral titers in the lung tissue and similar levels of viral shedding during acute infection. However, low-volume intranasal exposure results in milder weight loss during acute infection while intragastric exposure leads to a diminished capacity to regain body weight following the period of acute illness. Further, we examined both sex and age differences in response to SARS-CoV-2 infection. Male hamsters, and to a greater extent older male hamsters, display an impaired capacity to recover from illness and a delay in viral clearance compared to females. Lastly, route of exposure, sex, and age were found to influence the nature of the host inflammatory cytokine response, but they had a minimal effect on both the quality and durability of the humoral immune response as well as the susceptibility of hamsters to SARS-CoV-2 re-infection. Together, these data indicate that the route of exposure, sex, and age have a meaningful impact SARS-CoV-2 pathogenesis in hamsters and that these variables should be considered when designing pre-clinical challenge studies.

## Introduction

Severe acute respiratory syndrome coronavirus 2 (SARS-CoV-2) is a positive-sense RNA virus which first emerged in Wuhan, China in December 2019^1,2^. It is a member of the *Betacoronavirus* genus and is related to the high consequence pathogens severe acute respiratory syndrome coronavirus (SARS-CoV) and Middle Eastern Respiratory Syndrome (MERS)-CoV. SARS-CoV-2, the causative agent of coronavirus disease 2019 (COVID-19), rapidly spread across the globe, infecting over 100 million people and killing over three million individuals. The COVID-19 pandemic is an unprecedented public health emergency with profound ongoing economic and societal consequences.

For the majority of people, SARS-CoV-2 infection results in asymptomatic^3-5^ or mild infection not requiring hospitalization. Moderate disease generally requires hospitalization with or without administration of exogenous oxygen, and severe disease is often defined by hospitalization and need for high flow oxygen or non-invasive or invasive ventilation may result in death^6^. Symptoms include fever, fatigue, muscle aches, anosmia, dysgeusia, gastrointestinal (GI) disfunction, and respiratory symptoms such as cough and difficulty breathing^7,8^.

Complications include pneumonia, acute respiratory distress syndrome (ARDS), multi-organ dysfunction, and/or thrombotic events. Severe COVID-19 is further associated with thrombocytopenia, lymphopenia, and an aberrant pro-inflammatory cytokine response^9,10^. Although patients in all age groups can develop critical illness, the severity of COVID-19 is associated with advanced age and comorbidities including cardiovascular disease, diabetes, and obesity ^7,8,11^. Notably, males appear to be at greater risk of progressing to severe disease, which has been proposed to be a result of sex-specific differences in the immune response to infection, including auto-antibodies to type I interferons^12-14^.

SARS-CoV and SARS-CoV-2 transmission is believed to occur primarily through the deposition of short-range respiratory droplets and aerosols or from fomites/contaminated surfaces onto mucous membranes of the eyes, nose, and mouth or by direct inhalation into the lungs^15-17^. Angiotensin-converting enzyme 2 (ACE2), a SARS-CoV-2 host cell receptor, and transmembrane protease serine 2 (TMPRSS2), a serine protease that facilitates viral entry, are expressed in multiple cell types, including ciliated airway epithelial cells (i.e. nasal, bronchial, and bronchiolar cells); goblet cells; alveolar type II pneumocytes; and enterocytes^18,19^. These cell types have been shown to be among those that are susceptible to SARS-CoV-2 infection^20-22^. Although SARS-CoV-2 is a respiratory virus, gastrointestinal symptoms are routinely observed in COVID-19 patients^23-25^. Prolonged viral RNA (vRNA) shedding in the feces^26^ and the detection of SARS-CoV-2 nucleocapsid within enterocytes in both animal models^27,28^ and humans^29^ suggests that like MERS^30^ the GI tract may serve as an alternate site of SARS-CoV-2 infection.

A concerted worldwide research effort has identified a number of animal models of SARS-CoV-2 infection that are essential tools for improving our understanding of disease and for evaluating candidate vaccines and therapeutics. Several nonhuman primate (NHP) models have been described, including rhesus macaques (*Macaca mulattai*)^31,32^, cynomolgus macaques (*M. fascicularis*)^33^, and African green monkeys (*Chlorocebus sp*) ^34^. Multiple small animal models of SARS-CoV-2 infection have also been described, including transgenic mice that express human ACE2 (hACE2)^35,36^, conventional laboratory mice strains that are infected with a mouse-adapted SARS-CoV-2^37,38^, and several additional species that are naturally susceptible to infection with wildtype SARS-CoV-2, including tree shrews^39^, deer mice^40,41^, hamsters^27,42^, ferrets^28,43^, and domesticated cats^28,44^. Hamsters are a desirable model since upon intranasal exposure to SARS-CoV-2 they develop mild disease characterized by rapid breathing, weight loss, high viral loads in respiratory tissues, and extensive lung pathology^27,42^.

Given the value of the hamster model as a tool for better understanding SARS-CoV-2 pathogenesis and for evaluating prospective anti-viral countermeasures, we have sought to extend its utility by examining the impact on disease of route of inoculation, sex, and age on disease. Here we report that low-volume intranasal, high-volume intranasal, and intragastric routes of SARS-CoV-2 exposure result in a similar viral burden in the respiratory tract tissues and comparable levels of viral shedding; however, compared to high-volume intranasal exposure, low-volume intranasal exposure alone resulted in milder weight loss while intragastric exposure led to an impaired ability to regain body weight following the acute illness period. We further demonstrate a significant male bias in disease severity with male hamsters and to a greater extent older male hamsters displaying an impaired capacity to recover from illness. Remarkably, these differences in disease severity were not accompanied by differences in viral load in the examined tissues but did result in differential expression of inflammatory cytokines, perhaps suggesting a role for route of exposure, sex, or age-mediated immune factors in SARS-COV-2 pathogenesis. This study provides a comprehensive characterization of the Syrian golden hamster as a model for SARS-CoV-2 infection and demonstrates that route of virus administration, sex, and age are key variables influencing the course of disease. Appreciation of these variables is critical as efforts are undertaken to further our understanding of COVID-19 pathogenesis and to develop effective vaccines and therapeutics.

## Results

### Influence of route of exposure on SARS-CoV-2 pathogenesis in Syrian golden hamsters

To examine the influence of different routes of exposure on SARS-CoV-2 pathogenesis groups of five six-week old mixed male and female Syrian golden hamsters (*Mesocricetus auratus*) were exposed to 10^5^ TCID50 of SARS-CoV-2 by one of three routes of administration: intranasal low volume (20 µl, i.n.L), intranasal high volume (100 µl, i.n.H), and intragastric oral gavage (500 µl, i.g.). These routes of administration were selected to model upper airway (i.n.L), upper/lower airway (i.n.H), and oral/gastrointestinal (i.g.) routes of initial SARS-CoV-2 exposure (Fig. 1a).

**Figure 1:**
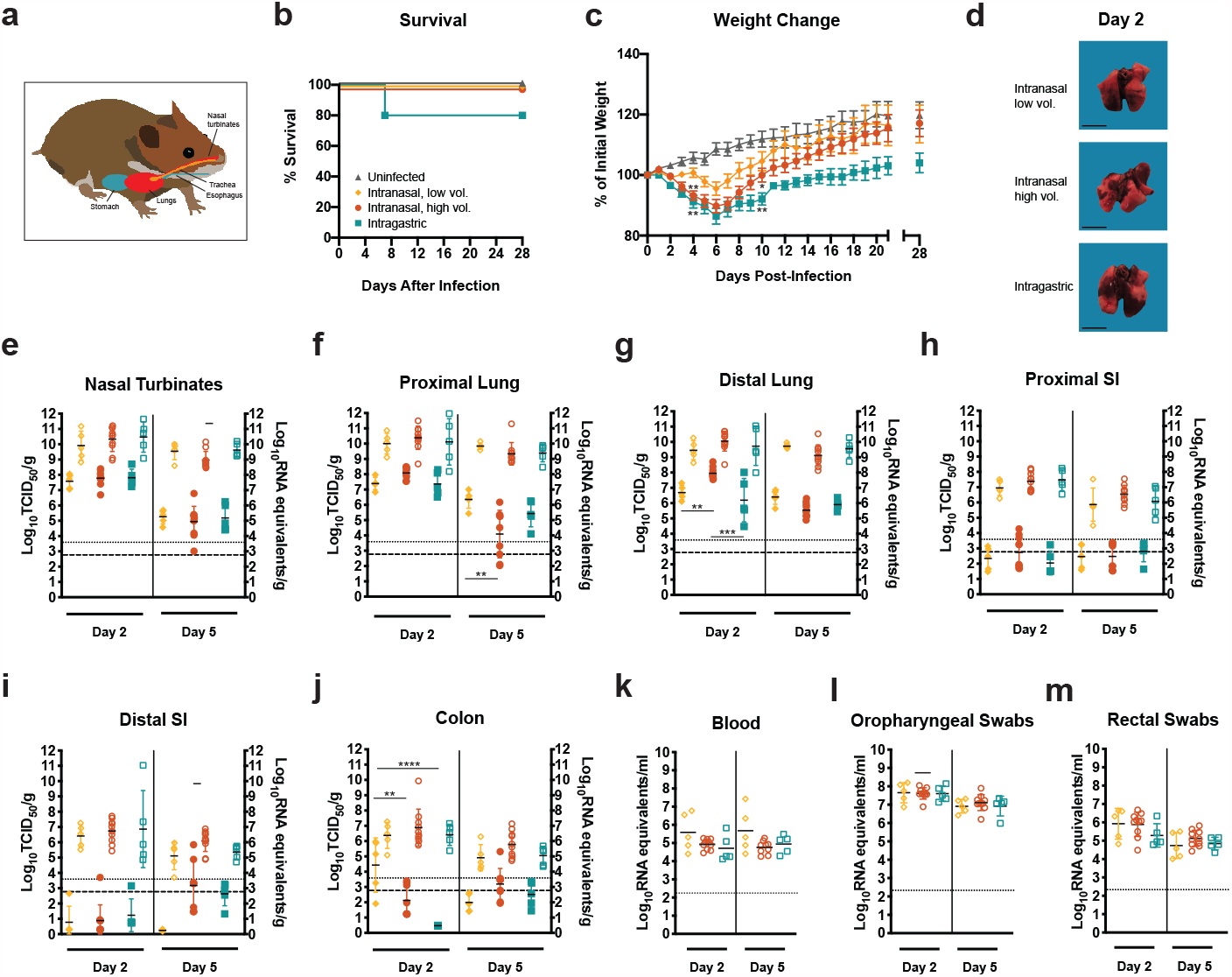
SARS-CoV-2 infection of adult golden Syrian hamsters by intranasal and intragastric routes of exposure. **a - m** Six-week-old mixed female and male Syrian golden hamsters (*Mesocricetus auratus*) were inoculated with 10^5^ TCID_50_ of SARS-CoV-2 by a low (20 µl) or high (100 µl) volume intranasal route of administration (orange diamonds or red circles, respectively) or an intragastric route of administration (teal squares) and compared to age-matched uninfected controls (grey triangles). Bars indicate means. Error bars indicate SEM (**c**) or SD (**e-m**). Dashed lines and dotted lines indicate the limit of detection for the TCID50 assay and qRT-PCR assay, respectively. **a**, Schematic depicting the routes of hamster inoculation. Kaplan-Meier curve depicting survival data (**b**) and weight data (**c**) over the course of 28 days after SARS-CoV-2 exposure. **d**, macroscopic images of SARS-CoV-2-infected hamster lungs. Scale bar = 1 cm. **e-m**, Infectious viral load (filled in shapes, left axis) and vRNA levels (empty shapes, right axis) in the (**e**) nasal turbinates, (**f**) proximal lung, (**g**) distal lung, (**h**) proximal small intestines, (**i**) distal small intestines, and (**j**) colon at the indicated days post-infection. Viral RNA levels in (**k**) blood, (**l**) oropharyngeal swabs and (**m**) rectal swab samples. Data were collected in a single experiment (n = 5).

Administration of SARS-CoV-2 by each of the three routes resulted in productive infection. SARS-CoV-2 exposed hamsters were monitored daily for clinical signs of illness for 28 days and showed no signs of illness other than weight loss and elevated respiratory rate and/or laboured breathing in some animals. Survival data is depicted as Kaplan-Meier survival curves (Fig. 1b). With the exception of a single hamster from the i.g.-exposed group that succumbed, all infected hamsters survived infection regardless of the route of SARS-CoV-2 exposure. Hamsters were weighed daily for 21 days post-infection (dpi) and again on 28 dpi (Fig. 1c). Mock-infected animals did not lose weight and gained weight throughout the course of the experiment. The majority of infected hamsters began to lose weight by 2 dpi, regardless of the route of SARS-CoV-2 exposure; however, at 4 dpi the i.n.H and i.g.-exposed hamsters showed significantly greater weight loss compared to the mock-infected control animals (One-way ANOVA, *P* = 0.0017 and 0.0011, respectively). Conversely, the hamsters exposed via the i.n.L route did not show significant weight loss (One-way ANOVA, *P* = 0.4250). At 4 dpi, the majority of the SARS-CoV-2-exposed hamsters belonging to the i.n.H and i.g. exposure groups showed weight loss ranging from 5-12%, and with the exception of one individual in the i.n.H group all hamsters had lost weight in both groups. In contrast, at the same time point three out of five of the i.n.L-exposed hamsters had gained weight, and the remaining two animals showed weight loss of only 1.1% and 4.2%. At 7 dpi the mean body weight had begun to rebound in all the virus-exposure groups; however, the increase in mean body weight of the i.g.-exposed hamsters lagged behind that of either i.n. exposed group. At 10 dpi, the mean weight of both i.n.-exposed groups had recovered to reach the initial starting weight, whereas the mean weight of the i.g. exposed hamsters remained significantly lower at ∼90% of the initial starting weight. Compared to the uninfected hamsters, the i.n.L-exposed hamsters weights were not significantly different (One-way ANOVA, *P* = 0.4243), whereas the i.n.H- and i.g.-exposed hamsters remained significantly lower (One-way ANOVA, *P* = 0.0499 and 0.038, respectively). Further, at 28 dpi the i.g.-exposed hamsters had re-gained less weight than the hamsters belonging to the uninfected and intranasal exposure groups, although these differences did not attain statistical significance. Sex differences were noted in the i.n.H group, which are discussed in the following section, below.

Additional groups of 6-week-old mixed sex hamsters were similarly infected with SARS-CoV-2 by i.n.L, i.n.H, and i.g. routes and were necropsied at 2 and 5 dpi (n = 5, 10, and 5, respectively at each time point). Macroscopic discoloration of portions of the lung tissue indicative of severe inflammation, often present in multiple lobes and sometimes encompassing more than half of the total lobe volume, was consistently observed in infected hamsters regardless of the route of exposure (Fig. 1d). The viral burden in the upper and lower respiratory tract, including the nasal turbinates, proximal lung, and distal lung was assessed (Fig.1e-g). At 2 dpi, high levels of infectious virus (∼ 10^7^ - 10^8^ TCID_50_/g) were detected in the nasal turbinates regardless of the route of exposure (Fig. 1e, solid shapes, left axis), and these values had declined similarly by several logs in all groups by 5 dpi. At 2 dpi, comparable levels of infectious virus (∼ 5×10^6^ – 5×10^8^ TCID_50_/g) were detected in the proximal lung regardless of the route of exposure (Fig. 1f, solid shapes, left axis). At 5 dpi the infectious virus in the proximal lung had declined by several logs in the i.n.H and i.g.-exposed hamsters and to a lesser extent in the i.n.L-exposed hamsters, resulting in a significantly greater infectious virus burden in the i.n.L versus i.n.H-exposed hamsters (One-way ANOVA, *P* = 0.0017). At 2 dpi, high levels of infectious virus were detected in the distal lung of all hamsters, regardless of the route of exposure (Fig. 1g, solid shapes, left axis); however, significantly more infectious virus was detected in the i.n.H exposed hamsters compared to the i.n.L and i.g.-exposed hamsters (One-way ANOVA, *P =* 0.0098 and 0.0002, respectively). At 5 dpi the level of infectious virus in the distal lung had significantly declined in the i.n.H.-exposed group (One-way ANOVA *P* < 0.0001), whereas the level of infectious virus in the i.n.L and i.g.-exposed hamsters remained similar to the 2 dpi level. At 2 dpi, high levels of viral RNA (vRNA) were detected in the nasal turbinates, proximal lung, and distal lung (vRNA; ∼ 10^8^ –10^12^ genome equivalents/g), regardless of the route of exposure and remained at a similar level at 5 dpi, having declined to a much lesser extent than the level of infectious virus (Fig. 1e-g, empty shapes, right axes). It is interesting to note that the i.g.-exposed hamsters displayed prolonged illness compared to the i.n.H-exposed hamsters group despite the presence of less infectious virus in the distal lung region at 2 dpi.

The infectious virus burden was also assessed in the digestive tract, including the small intestines (proximal and distal) and colon (Fig. 1h-j, solid shapes, left axes). Regardless of the route of infection, infectious virus in the GI tract tissues was below or close to the limit of detection, with the exception of some animals exposed by either i.n. route that had moderate levels in the colon (i.n.L route group, 2 dpi) or the distal small intestine and colon (i.n.H group, 5 dpi) (Fig. i-j). At 2 dpi, there was significantly more infectious virus present in the colon of the i.n.L-exposed hamsters compared to both the i.n.H and i.g.-exposed hamsters (One-way ANOVA *P =* 0.0029 and <0.0001, respectively). At 2 dpi moderate levels of vRNA (ranging from ∼ 2×10^5^ – 2×10^8^ genome copies/g) were detected in the proximal intestine, distal intestine, and colon, regardless of the route of infection (Fig. 1h-j, empty shapes, right axes), and these values had declined by 5 dpi. Despite reports that SARS-CoV-2 can infect small intestine enterocytes in vitro^21,22^, we found that a high titer SARS-CoV-2 challenge administered directly to the digestive tract failed to result in an increased viral burden in the small intestines or colon compared to i.n. inoculation.

Persistent SARS-CoV-2 RNAemia in humans has been linked to greater inflammation and more severe disease^45^. At both 2 and 5 dpi, vRNA was detected in the blood of all hamsters, regardless of the route of exposure (Fig. 1k). Low levels of infectious virus were detected in the blood at 2 dpi in a small number of samples (data not shown).

Lastly, to monitor viral shedding, oropharyngeal and rectal swab samples were collected prior to euthanization at 2 and 5 dpi (Fig. 1l-m). Moderate vRNA levels (ranging from ∼ 10^7^ – 10^8^ genome copies/ml in oropharyngeal swabs and 10^4.5^-10^6.5^ genome copies/ml in rectal swabs) were detected at 2 dpi in all groups, and vRNA levels detected in either swab sample type had declined by approximately one log at 5 dpi. The shedding of vRNA did not differ statistically between the exposure groups. Direct contact between infected and naïve hamsters has been shown to result in SARS-CoV-2 transmission^27,42^. The observed shedding data suggests that there may be similar shedding/transmission potential regardless of the route of exposure of an infected donor hamster. Notably, these vRNA shedding data imply that hamsters exposed by the i.n.L route that follow a milder disease course as evidenced by limited weight loss would be expected to have a similar capacity to transmit virus as the more overtly ill hamsters exposed by the i.n.H and i.g. routes.

### Influence of sex and age on SARS-CoV-2 pathogenesis in golden Syrian hamsters

Differential disease outcomes between the sexes or different age groups in animal models of SARS-COV-infection could be an important consideration for the pre-clinical evaluation of vaccines, antivirals, and candidate therapeutics^46^. In order to explore the contribution of sex to SARS-CoV-2 pathogenesis in hamsters we segregated the i.n.H-exposed hamster data, described above, by sex, resulting in two separate groups, 6-week-old males and 6-week-old females (n = 5 for each sex). An additional group of 22-week-old older male hamsters (n =5) were likewise infected with 10^5^ TCID^50^ of SARS-CoV-2 via an i.n.H route, and similar analyses were carried out. Female, male, and older male hamsters were all productively infected, and no individuals succumbed to disease (Fig. 2a). Mock infected animals did not lose weight and gained weight throughout the course of the experiment (Fig. 1b). Weight loss in all three infected hamster groups began on 2 dpi and tracked similarly until 5 dpi when the mean weights began to rebound at a faster rate in the females compared to the males and older males (Fig 2b). The SARS-CoV-2-infected female hamsters reached their pre-infection weights at 9 dpi and ultimately reached the mean weight of the uninfected hamsters at 11 dpi. Conversely, the infected males and older males had significantly lower mean weights compared to the females from 9 dpi onwards (One-way ANOVA *P* = 0.0026 and 0.0020, respectively). Older males did not recover their initial body weights even by day 28. The macroscopic discoloration observed in large portions of the lung tissue in SARS-CoV-2-infected hamsters at 2 dpi varied greatly in individuals and did not appear to differ by sex or age (Fig. 2c).

**Figure 2:**
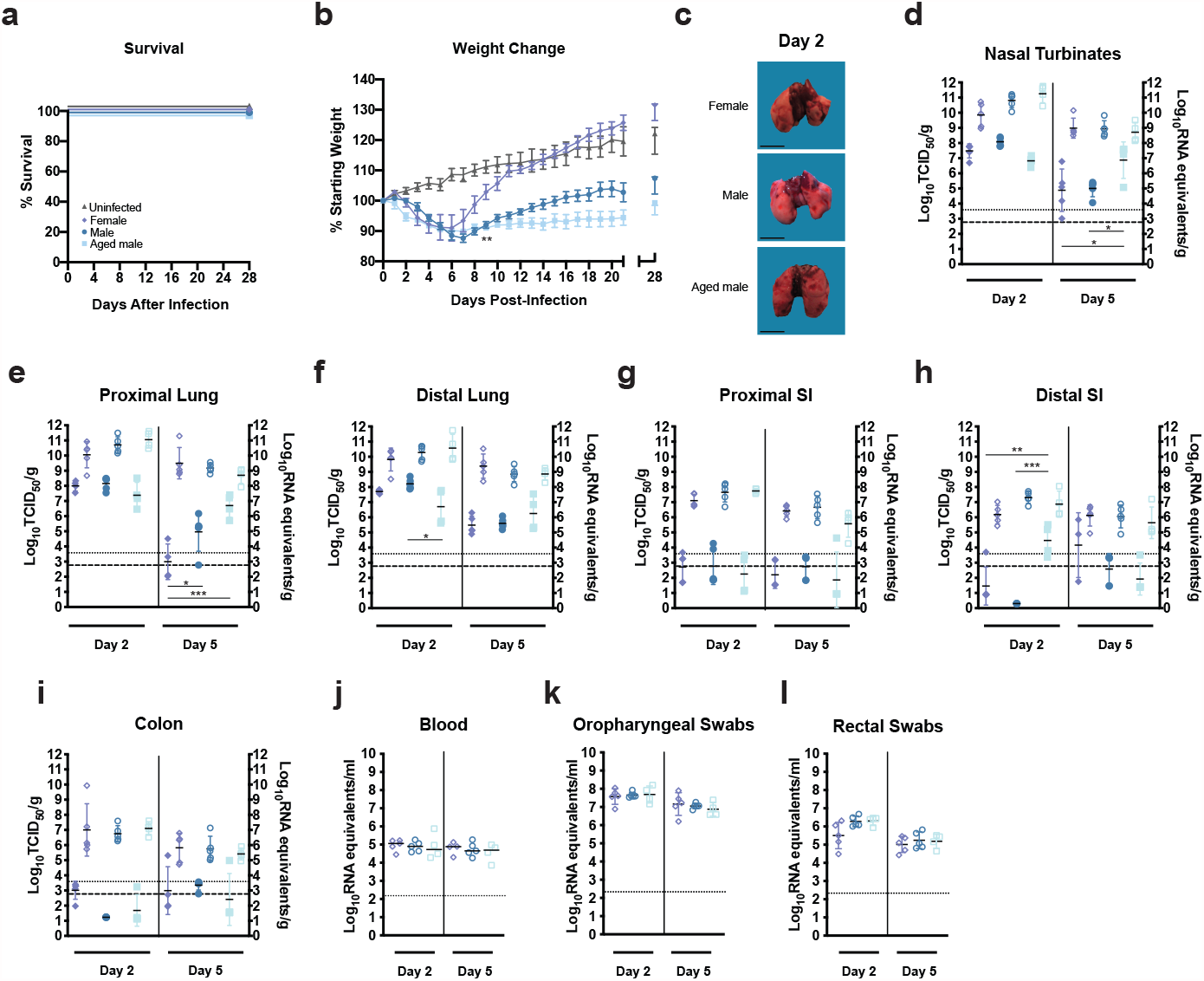
SARS-CoV-2 infection of female, male, and aged golden Syrian hamsters. **a - l** Six-week-old female (lavender diamonds), six week male (dark blue circles), or twenty-week-old male (light blue squares) hamsters (*Mesocricetus auratus*) were inoculated with 10^5^ TCID_50_ of SARS-CoV-2 by a high volume (100 µl) intranasal route (i.n.) of administration and compared to six-week old uninfected controls (grey triangles). Bars indicate means. Error bars indicate SEM (**b**) or SD (**d-l**). Dashed lines and dotted lines indicate the limit of detection for the TCID_50_ assay and qRT-PCR assay, respectively. **a-b**, Kaplan-Meier curve depicting survival data (**a**) and weight data (**b**) over the course of 28 days following SARS-CoV-2 exposure. **c**, macroscopic images of SARS-CoV-2-infected hamster lungs. **d-i**, Infectious viral load (filled in shapes, left axis) and vRNA levels (empty shapes, right axis) in the (**d**) nasal turbinates, (**e**) proximal lung, (**f**) distal lung, (**g**) proximal small intestines, (**h**) distal small intestines, and (**i**) colon at the indicated days post-infection. Viral RNA levels in (**j**) blood, (**k**) oropharyngeal swabs and (**l**) rectal swab samples. Data were collected in a single experiment (n = 5). The six week male and female data corresponds to the data presented for the high volume intranasal route of administration (Figure 1).

The infectious virus burden in the respiratory tract tissues for females, males, and older males was plotted (Fig. 2d-f, solid shapes, left axes). At 2 dpi the infectious virus burden in the respiratory tract tissues did not differ with the exception of the older males that showed a significantly lower viral burden in the distal lungs (One-way ANOVA *P* = 0.0364). At 5 dpi, the older males had significantly more infectious virus present in the nasal turbinates compared to females and males (One-way ANOVA *P* = 0.0175 and *P* = 0.0277, respectively). Both males and older males had significantly more infectious virus present in the proximal lungs, compared to females (One-way ANOVA *P* = 0.0347 and *P* < 0.0001, respectively). These data indicate that both sex and age differences occur in the hamster model of SARS-CoV-2 infection. These findings are consistent with the reported increased viral burden and delayed virus clearance reported in older mice compared to young mice exposed to mouse-adapted SARS-CoV-2^38^, but they are contrary to a prior report of SARS-CoV-2 infection in older hamsters, where no delay in viral clearance was observed^47^. The level of vRNA in the respiratory tract tissues did not differ show statistically significant differences among the three groups (Fig. 2d-f, empty shapes, right axes). The infectious virus burden in the digestive tract (proximal and distal small intestines and colon) was similar among the three groups (Fig. 2h-i, solid shapes, left axis), with the exception of the older males at 2 dpi that had significantly more infectious virus in the distal small intestine compared to both the females and young males (One-way ANOVA *P* = 0.0096 and 0.0003, respectively). The levels of vRNA in the digestive tract did not show statistically significant differences among the three groups (Fig. 2 g-i, empty shapes, right axes). At both 2 and 5 dpi, vRNA was detected in the blood of all hamsters, and the levels did not differ among the three groups (Fig. 2j). The observed differences in weight between male and female mean body weight throughout the course of illness does not appear to be due to large differences in viral burden in the tissues of the respiratory tract or GI tract, and therefore may be due to the host response to infection driving immune-mediated pathology resulting in prolonged sickness.

Moderate vRNA levels ranging from ∼ 10^7^-10^8^ genome copies and 10^4.5^-10^6.5^ genome copies/ml were detected at 2 dpi in oropharyngeal and rectal swabs, respectively, and the vRNA levels did not differ among the different groups (Fig. 2k-l). These data suggests that regardless of the differential disease presentation in female, male, and older male hamsters there may be similar shedding/transmission potential during acute illness irrespective of the sex or age.

The lesions observed in the lung tissue of individual hamsters at 5 dpi were variable and ranged in severity and progression within each inoculation route group. Although the entire spectrum of lesions could be observed within each group there were some differences between groups in the frequency in which specific lesions were observed. In some animals, lung lesions were more typical of earlier infection and characterized by patchy bronchointerstitial pneumonia with necrosis of alveolar and bronchiolar epithelial cells. Interstitial infiltrates consisted primarily of neutrophils and macrophages. There was alveolar edema/hemorrhage as well as perivascular edema. Other animals had lesions that were more typical of later infection with severe interstitial pneumonia (lymphohistiocytosis) leading to loss of alveolar spaces. There was extensive type II pneumocyte hyperplasia with characteristic tombstoning, peribronchitis, hyperplasia of bronchiolar epithelial cells, perivasculitis, presence of multinucleated syncytial cells and occasionally vasculitis or fibrosis. Representative histology images from each group are presented (Fig. 3a, upper panel). In general, more animals in the i.n.H group had lesions of patchy bronchointerstial pneumonia compared to the i.g. group, where lesions were characterized by severe, widespread lymphohistiocytic interstitial pneumonia. In the i.n.L group, there were roughly equal numbers of each type of lesion. Due to the limited small sample size additional experiments would be required to confirm these putative differences. In situ hybridization (ISH) staining showed the presence of vRNA, primarily detected in bronchiolar epithelial cells, that was highly variable between individual animals and/or samples (Fig. 3a, lower panels). Lesions were largely absent from the nasal turbinates (data not shown) although vRNA was detected in some epithelial cells (Extended Data Fig. 1).

**Figure 3:**
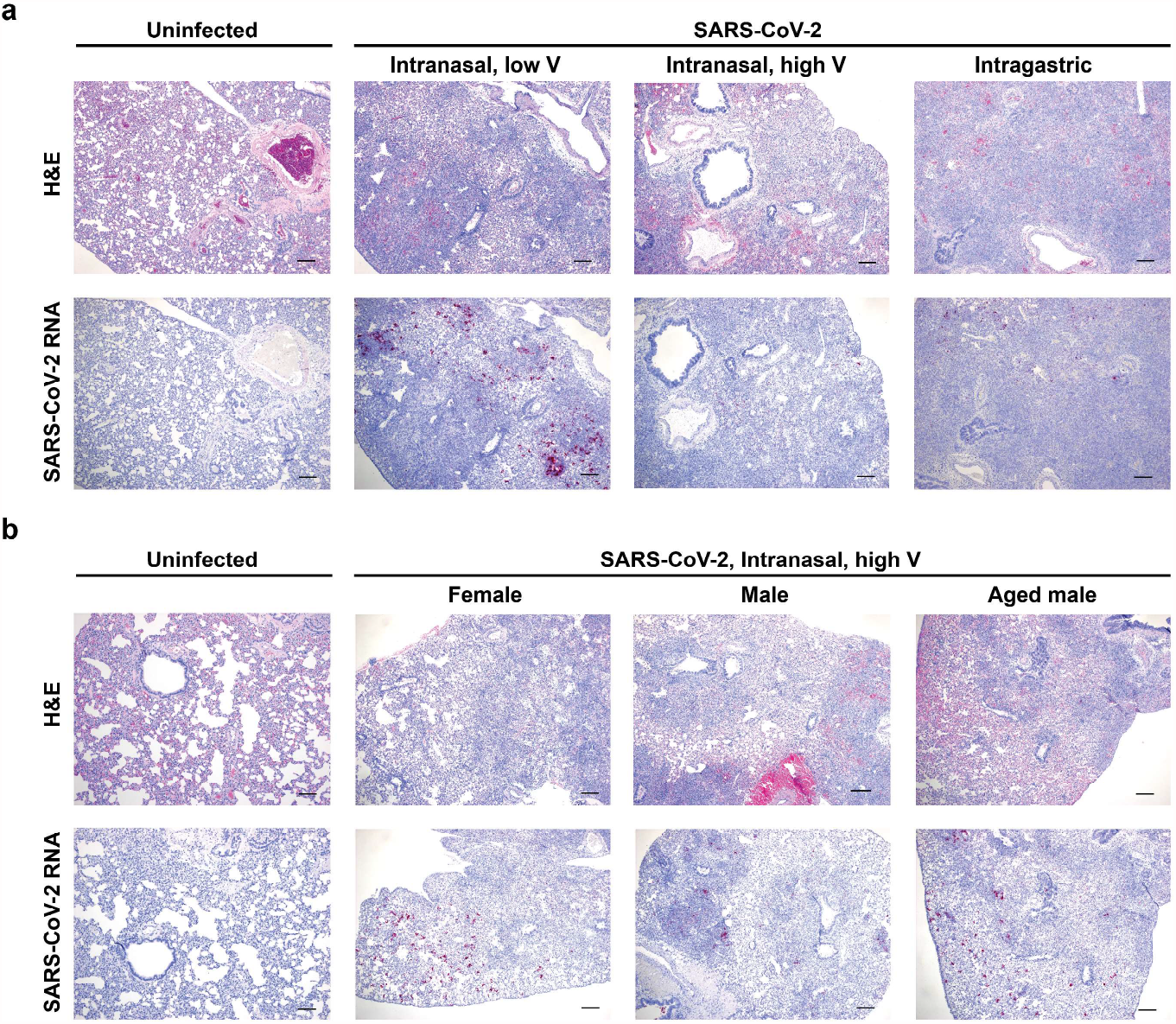
Histopathology and virus distribution in the lungs. Hematoxylin/eosin (H&E) staining (**a, b**, upper panels) and in situ hybridization (ISH) with antisense probes that detect the SARS-CoV-2 genome/mRNA (**a, b**, lower panels) on lung tissue of uninfected and SARS-CoV-2-infected hamsters at 5 dpi. **a**, six-week-old mixed female and male golden hamsters (*Mesocricetus auratus*) were inoculated with 10^5^ TCID_50_ of SARS-CoV-2 by a low (20 µl) or high (100 µl) volume intranasal route of administration or an intragastric route of administration and compared to age-matched uninfected controls. **b**, six-week old female, six-week old male, or twenty-week old male hamsters were inoculated with 10^5^ TCID_50_ of SARS-CoV-2 by a high volume (100 µl) intranasal route of administration and compared to six-week old uninfected controls. Positive detection of viral genomic RNA/mRNA is indicated by magenta staining. The magnification is 5x for H&E and ISH. Scale bars = 200 µm.

Similarly, a comparison of males and females showed a wide variability in lesion severity in individual animals within each group. Overall, the spectrum of lesions observed was similar in both females and males, and no differences in lesion severity were observed (Fig. 3b, upper panel). Similarly, no differences in the nature or severity of lung lesions were observed in the younger compared to the older males. Due to the wide variability between individual animals, a larger sample size is needed to detect subtle differences between the sex groups and age groups. These observations are consistent with the variable lung pathology observed in SARS-CoV-2-infected cynomolgus macaques^33^. ISH staining showed similar levels of vRNA in females, males, and older males (Fig. 3b, lower panels).

Blood biochemistry and hematological parameters were assessed for both mock-(both sexes, n = 6) and SARS-CoV-2-infected hamsters belonging to the various experimental groups described above, including the i.n.L, i.n.H, and i.g. exposure groups (both sexes, n = 5, 10, and 5 for the exposure groups, respectively); the i.n.H group broken down by sex (n = 5 of each sex); and the i.n.H-exposed older males (n = 4) (Extended Data Fig. 2 a-g). At 5 dpi all the groups had similar counts of white blood cells (Extended Data Fig. 2a), lymphocytes (Extended Data Fig. 2b), and neutrophils (Extended Data Fig. 2c); however, the neutrophil-to lymphocyte ratio (NLR), a clinical metric that has been found to be elevated in human patients during severe COVID-19^48^, was significantly elevated in the i.n.H (both sexes combined) and i.n.H-exposed male hamsters compared to the uninfected control hamsters (One-way ANOVA *P* = 0.0052 0.0014) (Extended Data Fig. 2d). At 2 dpi alanine aminotransferase (ALT) values were not significantly elevated in any group, suggesting that liver damage had not occurred (Extended Data Fig. 2e), whereas blood albumin (ALB) was reduced in the i.n.L, i.n.H, i.g., and female i.n.H-exposed compared to uninfected hamsters (Extended Data Fig. 2f, One-way ANOVA *P* = 0.0009, 0.0048, 0.0002, 0.0005, respectively), recapitulating the ALB reductions observed in severe COVID-19 patients^49^. Blood urea nitrogen (BUN) was elevated in all groups, suggestive of potential kidney disfunction or pre-renal damage (dehydration), regardless of route of infection, sex, or age (Extended Data Fig. 2g) although the presence of vRNA was not detected in the kidneys by ISH (data not shown). Additional serum biochemical values were unremarkable (data not shown).

### Influence of route of infection, sex, and age on the immune response to SARS-CoV-2 infection

With little variation in viral burden in the tissues observed between groups of animals despite notable differences in the course of clinical disease, we sought to determine whether route, sex, and/or age differentially impacts the immune response to SARS-CoV-2 infection. We examined the relative mRNA expression of a subset of inflammatory cytokines and immune response genes. Relative gene expression in the blood and lungs of IL-1β, IL-6, TNF-α, Mx-2, and STAT-2 at 2 dpi and IL-2, IL-4, IL-10, IFN-γ, and FoxP3 at 5 dpi was quantified for all of the experimental groups described above as compared to uninfected controls for baseline levels (Fig. 4a). The expression of inflammatory cytokines in the blood at 2 dpi was highly variable among individuals and did not differ significantly between any of the experimental groups (Fig. 4a, left panel). The route of virus administration impacted inflammatory cytokine expression in the lungs at 2 dpi (Fig. 4a, left middle). Overall, the mean of the examined cytokines at 2 dpi trended lower in the i.n.L group relative to the i.n.H group, and was lower still in the i.g. group, although these differences did not always attain statistical significance. Expression of IL-1β and IL-6 mRNA was significantly lower in the i.n.L-versus the i.n.H-exposed hamsters (Multiple t test *P* = 0.025 and 0.025, respectively). Further, lung expression of IL-1β, IL-6, TNF-α, Mx-2, and STAT-2 mRNA was significantly less in the i.g.-versus the i.n.H-exposed hamsters (Multiple t test *P* = 0.033, 0.014, 0.017, 0.0011, and 0.014, respectively). Sex and age were not found to have a significant effect on inflammatory cytokine expression in the lungs at this time point. In the older males transcription of IL-1β and IL-6 trended higher and transcription of STAT-2 trended lower relative to the young males, although these differences did not attain statistical significance. Differential expression of inflammatory cytokines became more pronounced between the infection groups at 5 dpi when we examined the expression of adaptive immune response genes in the blood and lungs of infected hamsters relative to uninfected controls. Inflammatory cytokine expression in the blood at 5 dpi was similar in the two i.n.-exposed groups (i.n.L and in.H), with only small differences observed that did not attain statistical significance. Cytokine expression trended lower in the i.g.-relative to the i.n.-exposed groups, and the i.g. group had significantly lower IL-10, IFNγ, and Fbp3 mRNA expression relative to the i.n.L group (Multiple t test *P* = 0.006, 0.012, 0.033, respectively). At 5 dpi, the young and older males showed similar cytokine expression in the blood, whereas cytokine expression in the females consistently trended higher, with females showing significantly elevated expression of IL-2, IL-4, and IL-10 relative to the young males (Multiple t test *P* = 0.034, 0.002, 0.029, respectively) (Fig 4a, middle right panel). Finally, consistent with overall cytokine expression observed in the blood and lung samples, IL-2 and IL-4 trended lower in the i.g. group relative to both i.n. groups, although these differences did not attain statistical significance. Inflammatory cytokine expression in the lungs at 5 dpi did not show statistically significant differences between the i.g.- and i.n.H-exposed hamsters; however, the i.n.L-exposed hamsters had elevated IL-10 and IFN-γ relative to the i.n.H-exposed hamsters (Multiple t test *P* = 0.007 and 0.013, respectively) and elevated IL-4, IL-10, and IFN-γ relative to the i.g.-exposed hamsters (Multiple t test *P* = 0.04, 0.01. 0.01, respectively) (Fig. 4a, right panel). At this time point inflammatory cytokine expression was similar in age-matched males and females although female hamsters had significantly reduced expression of IL-10 in the lungs compared to males (Multiple t test *P* = 0.029). In the older males inflammatory cytokine expression in the lungs consistently trended higher relative to the young males, although these differences did not attain statistical significance. Taken together these data suggest that differences in expression of inflammatory cytokine and immune response genes may drive differences in the SARS-CoV-2 disease course in hamsters.

**Figure 4:**
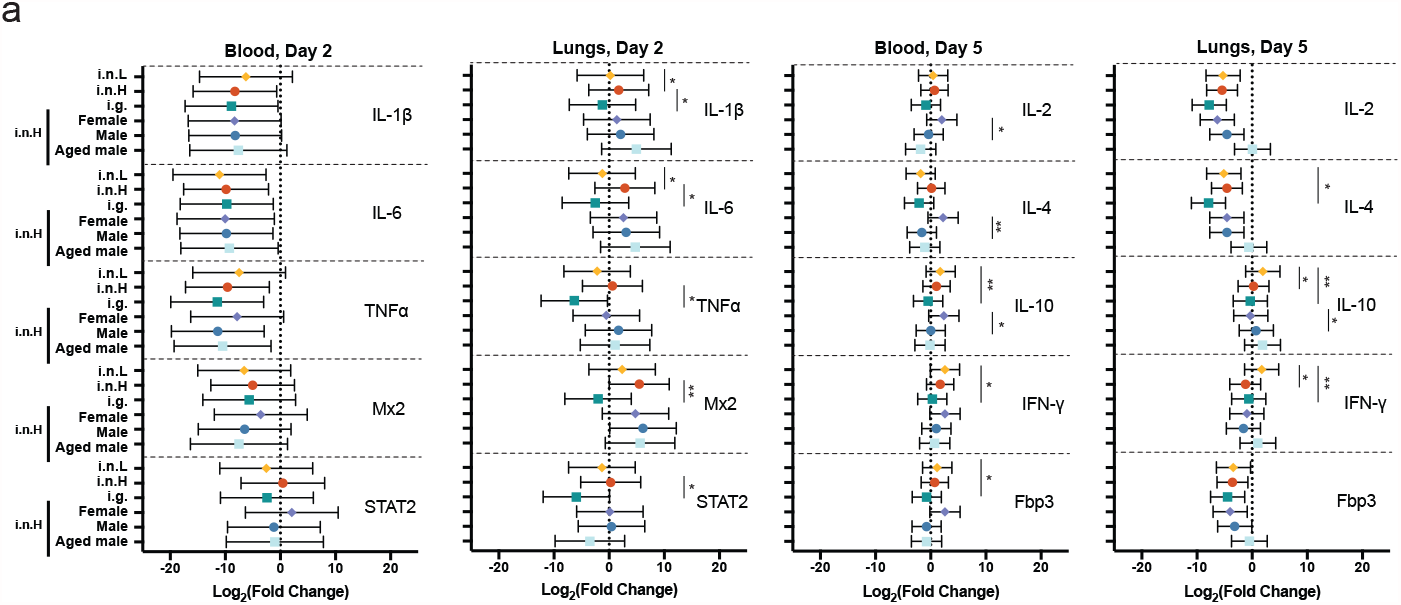
Hamster host response to SARS-CoV-2 infection. **a**, Six-week-old male and female hamsters (*Mesocricetus auratus*) were inoculated with 10^5^ TCID50 of SARS-CoV-2 by a low volume intranasal (orange diamonds), high volume intranasal (red circles) or intragastric (teal squares) route of administration. The high volume intranasal group data is further broken down by sex (female, lavender; male, dark blue) and compared with an additional group of 20 week old males exposed to 10^5^ TCID_50_ of SARS-CoV-2 by a high volume intranasal route of exposure (light blue squares). Cytokine gene expression was measured for IL-1β, IL-6, TNFα, Mx2, and STAT2 in the blood (left panel) and lungs (middle left panel) at 2 dpi and IL-2, IL-4, IL-10, IFN γ, and Fbp3 in the blood (right middle panel) and lungs (right panel) at 5 dpi and displayed relative to age-matched mock-infected animals. Gene expression was normalized using RPL18 as a control. Bars indicate mean, error bars indicate 95% confidence interval (a). * = *P* < 0.05, ** *P* = 0.01, ns = *P* > 0.05; unpaired student t test.

The humoral response generated following SARS-CoV-2 infection was evaluated in all the experimental groups at 21dpi (Extended Data Fig. 3) and 81 dpi (Fig. 5a-b). All SARS-CoV-2-exposed hamsters had detectable serum IgG titers against spike antigen as assessed by ELISA at 21 dpi with endpoint titers ranging from 1600 to 6400, the limit of the assay, that did not differ regardless of route of exposure, sex, or age (Extended Data Fig. 3a). At 81 dpi IgG titers ranged from 6400 to 102400, and there was a high level of variation within each group (Fig. 5a). While the i.g.-exposed hamsters had lower IgG titers relative to the i.n.-exposed hamsters, and the older males had lower IgG titers relative to the young males these differences did not attain statistical significance. Similarly, neutralizing antibodies were detected by plaque reduction neutralization test (PRNT^90^) at 21 dpi in all infected hamsters ranging from 1:80 to 1:1280, and no significant differences were observed regardless of route of exposure, sex, and age (Extended Data Fig. 3b). The neutralizing antibody levels stayed at approximately the same level when they were assessed at 81 dpi (Fig. 5b). Older males generally had lower levels of neutralizing antibodies although these differences did not attain statistical significance.

**Figure 5:**
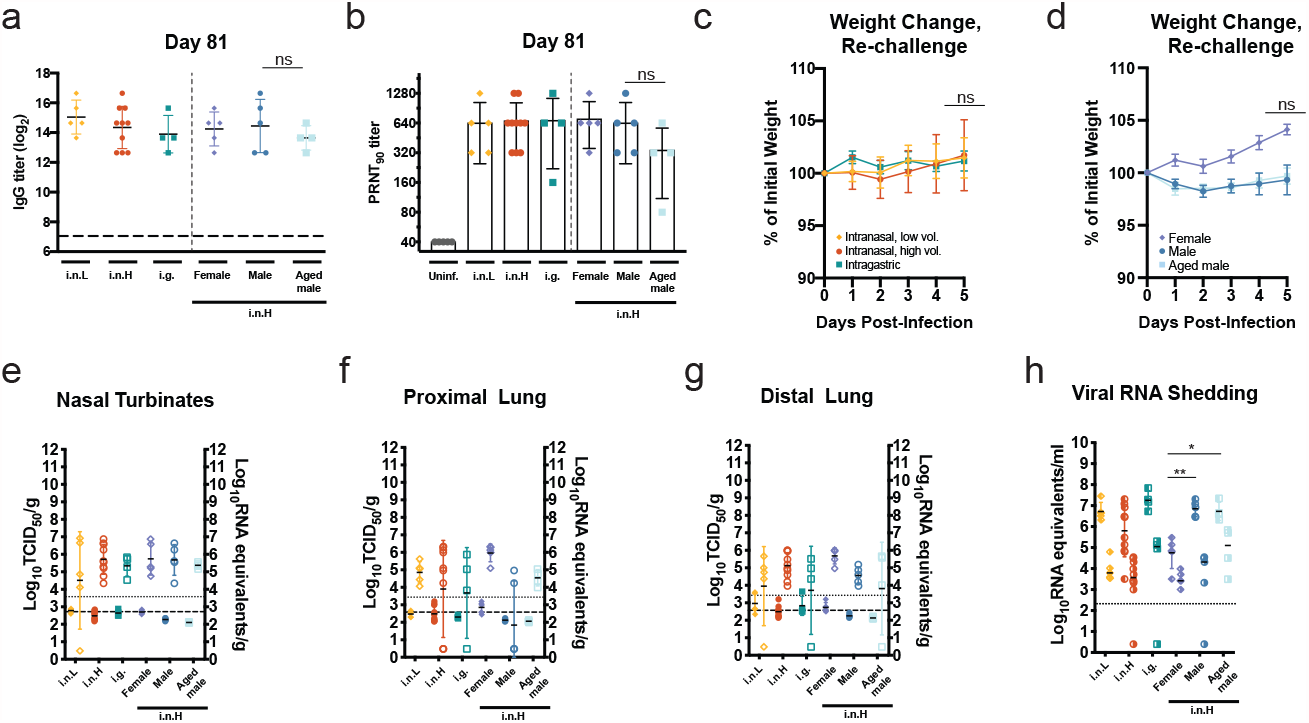
Durability of humoral immune response and susceptibility to re-challenge. **a-h**, Six-week-old male and female hamsters (*Mesocricetus auratus*) were inoculated with 10^5^ TCID_50_ of SARS-CoV-2 by a low volume intranasal (orange diamonds), high volume intranasal (red circles) or intragastric (teal squares) route of administration. The high volume intranasal group data is further broken down by sex (female, lavender; male, dark blue) and compared with an additional group of 20 week old males exposed to 10^5^ TCID50 of SARS-CoV-2 by a high volume intranasal route of exposure (light blue squares). **a**, IgG antibody response against SARS-CoV-2 spike antigen were assessed by ELISA using serum collected at 81 dpi. **b**, Neutralizing antibody against SARS-CoV-2 was measured by PRNT_90_ using serum collected at 81 dpi. **c-h**, Five hamsters belonging to the each of the challenge groups described above were re-challenged at 81 dpi with 1×10^5^ TCID_50_ of the same strain of SARS-CoV-2 by a high volume intranasal route of administration. **c-d**, Weights were obtained daily until 5 dpi for re-challenged hamsters initially exposed by an i.n.L, i.n.H, or i.g. route (c) or female, male, and older hamsters (d). **e-g**, Infectious virus (left axis, shaded) and viral RNA levels (right axis, empty) were measured in the nasal turbinates (**e**), proximal lung (**f**), and distal lung (**g**). **h**, Viral RNA shedding detected in oropharyngeal (left shaded) and rectal swabs samples (right shaded). Dashed horizontal lines indicate the limit of detection. The limits of detection are indicated with dashed lines (ELISA and TCID_50_ assay) and dotted lines (qRT-PCR assay). Bars indicate mean, error bars indicate SD (a-h). * = *P* < 0.05, ** *P* = 0.01, ns = *P* > 0.05; unpaired student t test.

At 81 dpi the infected hamsters belonging to each group were re-challenged by an i.n.H route with 1×10^5^ TCID50 of the same strain of SARS-CoV-2, and swab and tissues samples were obtained at 5 days after re-infection. Upon re-challenge some hamsters were observed to lose a small amount of weight (Fig. 5c). While the route of initial exposure and age did not appear to affect post re-challenge weight loss up to 5 dpi, female hamsters gained a slight amount of weight following re-challenge, whereas the males and older males lost a small amount of weight, however, these differences did not attain statistical significance (Fig. 5c-d). The viral burden in the respiratory tract tissues of the re-challenged hamsters were also assessed at 5 dpi (Fig. 5e-g). The amount of infectious virus and vRNA in the nasal turbinates (Fig. 5e), proximal lung (Fig. 5f), and distal lung (Fig. 5g) were several logs lower than the levels observed during the acute period following the initial SARS-CoV-2 exposure (Fig 5e-g versus Fig. 1e-g and 2d-f). The level of infectious virus in the respiratory tract tissues was consistently near or below the limit of detection of the assay. An additional group of i.n.H-exposed male hamsters that were re-challenged 133 (n = 5) or 140 (n = 14) showed very mild weight loss in some hamsters although these weight changes did not attain statistical significance (Extended Data Fig. 4). At 5 days after re-challenge (145 dpi), only two of fourteen hamsters had infectious virus near the limit of detection in the nasal turbinates, whereas all other organs tested (proximal lung, distal lung, and small intestines) contained no detectable infectious virus (Extended Data Fig. 4). This apparent protection correlated with a robust antibody response that was detectable at high titers at 140 dpi (Extended Data Fig. 4), suggesting that protective antibody and potentially protective memory responses can persist.

Interestingly, the levels of shed vRNA in oropharyngeal and rectal swab samples following the 81 dpi re-challenge were comparable to the levels observed during the acute period following the initial SARS-CoV-2 exposure (Fig. 1l-m and 2k-l versus Fig 5h). Re-challenged males and older males shed significantly more vRNA in oropharyngeal swab samples compared to females (*P* = 0.0053 and 0.0159, respectively) (Fig. 5h). The i.g.-exposed and re-challenged hamsters shed more vRNA in rectal swabs compared to re-challenged i.n.-exposed hamsters, and re-challenged males and older males had more vRNA in rectal swabs compared to re-challenged females; however, these differences did not attain statistical significance. These data suggest that route of exposure, sex, and age do not have a meaningful impact on the protective efficacy of the host immune response resulting from a natural exposure against a homologous SARS-CoV-2 re-exposure at up to 81 days following the initial exposure; however, these data suggest that males and older males shed more vRNA in oropharyngeal swabs than females. At 5 dpi following re-challenge the IgG titer had increased in most of the hamsters (Extended Data Fig. 4).

## Discussion

The unified global push to rapidly develop antivirals, experimental therapeutics, and vaccines that are capable of mitigating COVID-19 severity in infected individuals and/or virus transmission relies on animal models of SARS-CoV-2 infection that recapitulate with reasonable fidelity the key disease features of COVID-19. Here, we build upon previous reports that Syrian hamsters are a suitable animal model for SARS-CoV-2 infection ^27,42,47^ and describe the contribution of route of exposure, sex, and age to SARS-CoV-2 pathogenesis. We report that the route of inoculation influences the course of disease in golden Syrian hamsters. Compared to high-volume intranasal exposure, a low-volume intranasal exposure resulted in milder weight loss while intragastric exposure resulted in slower recovery from weight loss following the period of acute illness. Further, the route of SARS-CoV-2 exposure influenced the viral burden in a subset of respiratory tract and GI tract tissues, while the viral shedding and humoral immune responses were comparable. We also report that male hamsters, and to a much greater extent older male hamsters, demonstrated a reduced capacity to recover from illness compared to females showing significant differences in disease course related to sex and age. Males and older males also showed delayed viral clearance from the respiratory tract tissues compared to females; however, similar humoral immune responses were identified in females, males, and older males up to 81 dpi. We further identified variable expression of inflammatory cytokines in the blood and lungs with route of exposure, sex, and age, and these differences may point to a putative mechanistic explanation for the observed differences related to clinical presentation of disease in the various groups. Lastly we found that the route of initial exposure, sex, and age did not significantly influence the course of disease following a homologous SARS-CoV-2 re-challenge at 81 dpi although we found that females shed significantly less vRNA upon re-challenge than males and older males.

Studies describing SARS-CoV-2 pathogenesis in golden hamsters have employed various routes of virus administration, including intranasal with inhalation^27,42^, combined ocular and intranasal^47^, and oral administration^50^. Variable disease outcomes in response to different routes of inoculation, including intragastric inoculation, have been described for several high consequence respiratory viruses, including H5N1 influenza A virus^51^, Nipah virus (NiV)^52^, and MERS-CoV^30^; and respiratory tract exposure has been shown to result in productive infection of the gastrointestinal tract following viremia with the enterovirus, porcine epidemic diarrhea virus (PEDV)^53^. The previously reported intranasal administrations of SARS-CoV-2 to hamsters employed an inoculum volume ranging from 80 to 100 µl^27,42^. The volume of the inoculum delivered intranasally has been shown to dictate the efficiency of lower respiratory tract delivery, with a smaller volume more restricted to the upper respiratory tract ^54^. We sought to compare SARS-CoV-2 pathogenesis following a low (20 µl, i.n.L) or high (100 µl, i.n.H) inoculum volume to primarily limit the initial infectious dose to the upper respiratory tract or combined upper/lower respiratory tract, respectively. Further, COVID-19 patients often present with GI symptoms, SARS-CoV-2 vRNA is often be detected in the feces^23-26^, and SARS-CoV-2 nucleocapsid protein has been detected within enterocytes in both animal models^27,28^ and humans^29^. We, therefore, sought to further evaluate SARS-CoV-2 pathogenesis following an intragastric route of exposure. The described data demonstrates that an i.n.L inoculation resulted in a milder course of illness with reduced weight loss compared to the standard i.n.H inoculation, whereas an i.g. inoculation also resulted in efficient infection in the respiratory tract but with much slower recovery to the original body weight following acute illness. This is in contrast to the mild disease reported following SARS-CoV-2 infection of hamsters by an oral route^50^. Further, the only animal to succumb to infection had been exposed by an i.g. route. The i.n.L route of exposure may serve as a useful model for the asymptomatic SARS-CoV-2 infection with viral shedding that is commonly observed in humans^3,4^. While the acidic environment of the digestive tract is expected to inactivate virus SARS-CoV-2 within a matter of hours^21,50^ human enterocytes express both ACE2 and TMPRSS2^19^ and support virus replication in vitro^21,22^. Further, we observed rare and sporadic foci of vRNA in the GI tract by ISH (data not shown) as well as infectious virus in some GI tissue samples (Fig. 1 and 2). Taken together, these data suggest that a low level of viral replication may be occurring in the GI tract. As with MERS infection^30^, the GI tract may serve as a site of extrapulmonary SARS-CoV-2 replication and may contribute to viral pathogenesis or the nature of the immune response generated in response to infection. The variable levels of infectious virus detected in the digestive tract could be related in part to the feed state of the hamsters at the time of infection since simulated gastric and/or colonic fluids (particularly in the fasted state) have been shown to inactivate both MERS^30^ and SARS-CoV-2^21^. A limitation of the routes of administration described in this work is that i.n. adminstration results in some inoculum being swallowed and entering the gastrointestinal tract^54^.

Even with low volume i.n. administration, there may be a small amount of virus that become deposited within the lower respiratory tract shortly after infection. With the i.g. instillation by oral gavage some inoculum is also expected to be introduced to the respiratory tract^52^ or oral cavity when the feeding tube is inserted or withdrawn. We questioned whether the differential pathogenesis observed with an i.g. route versus an high-volume intranasal route administration was due to the alimentary tract exposure or rather a small viral dose seeding the airway during gavage; however, a recent report describes that low dose SARS-CoV-2 inoculation of the hamster airway results in diminished pathogenesis^55^. Given that the i.n.L inoculation may also be expected to result in a low dose delivery of virus to the lower respiratory tract, the milder clinical signs of illness that we observed are consistent with the reduced pathogenesis reported by others upon low dose exposure. Given our findings that all routes of infection resulted in similar timing and extent of virus replication in the respiratory and digestive tract tissues (Fig. 1), it is possible that disease outcome is more directly related to the nature of the immune responses that are established at the initial site of greatest virus exposure and that this has a significant impact on the cytokine expression at subsequent sites of active virus replication. Further, the high viral burden in the respiratory tract with viral shedding and reduced disease severity observed the i.n.L-exposed hamsters appears to recapitulate the asymptomatic disease state observed in most humans^4^. In contrast to a previous study^56^, we did not observe an age-dependent differences is lung pathology (Fig. 3).

Epidemiological and clinical analyses have identified several risk factors that are associated with an increased incidence of severe and and/or fatal COVID-19, including hypertension, cardiac, or pulmonary disease^8,57^, male sex ^58^, and advanced age ^10,57,58^; however, insight into how these factors contribute to SARS-CoV-2 disease progression in small animal models has only begun to be examined^47,56^. Several studies have reported that SARS-CoV-2 infection in male patients results in a greater risk of severe disease outcome than in females, potentially due to differences in the host response to infection^12,13,59,60^. We report that compared to i.n.H-infected females, males regained their body weight at a slower rate following the period of acute illness. We observed a greater viral burden in the nasal turbinates and proximal lung tissues of young males and older males compared to young females. This is in agreement with sex differences observed in SARS-CoV-infected mice, where males were shown to have elevated viral titers in the respiratory tract^61^. In that work the sex differences were attributed to the influence of sex hormones on the innate immune response rather than the adaptive response^61^, and it will be interesting to determine if this is consistent in the hamster model of SARS-CoV-2 infection. It has been shown that females can often mount a stronger inflammatory response in response to viral infection and vaccination compared to males^62^. Representation of both sexes will therefore be crucial for pathogenesis studies and pre-clinical evaluation of vaccines against COVID-19. A correlation between advanced age and increased risk of severe disease and mortality associated with SARS-CoV-2 patients has been well established^7,8,57^. Greater disease severity has also been reported in both older murine and hamster models of SARS-CoV-2 infection^37,38,47,63^. Our observation that older hamsters show a delayed recovery of their body weight following i.n. SARS-CoV-2 infection is consistent with these reported data^47,56^.Our findings are also consistent with the reported increased viral burden and delayed viral clearance reported in older mice compared to young mice exposed to mouse-adapted SARS-CoV-2^38^, but are contrary to the reported absence of delayed viral clearance in SARS-CoV-2-infected older hamsters^47^.

Independent studies have identified elevated cytokine expression in the blood in severe human cases of COVID-19. Some of those that are thought to play a role in disease development include IL-1β, IL-6, TNF-α, IL-10, IL-2 and IFN-γ^10,64^. Our observation that hamster lungs at 2 dpi showed that mRNA expression levels for IL-1β and IL-6 were reduced in i.n.L-exposed hamsters and IL-1β, IL-6, TNF-α, Mx-2, and STAT-2 were reduced in i.g.-exposed hamsters relative to their i.n.H-exposed counterparts indicate that initial exposure of the virus by the i.n.L or i.g. route resulted in a less robust inflammatory response in the lungs. Additionally, IL4, IL10, and IFN-γ mRNA expression in the lungs were found to vary by route of infection at 5 dpi (Fig. 4). The differences in adaptive immune response gene expression observed between the i.n.L, i.n.H, and i.g. groups suggest that the initial site of virus exposure has an impact on the cytokine profile and likely the nature of the immune response developed in the lungs. This observation could have important implications for understanding immune-mediated pathology, anti-viral immunity, and vaccination strategies. It is reasonable to conclude that differences in the innate immune response, including the cell types present and differential expression and activation of pattern recognition receptor (PAMP) in various tissues, as well as draining lymph nodes can dictate the nature of the adaptive immune response in response to SARS-CoV-2 infection^65-67^.

Interestingly, while significant differences in the expression levels of cytokines between young and older hamsters were not detected, older hamsters had consistently higher expression of several examined cytokines in the lungs, consistent with a study describing a stronger inflammatory response in the lungs of aged SARS-CoV-infected NHPs^68^. There remains a lack of developed, commercially available reagents for examining specific aspects of immunity in hamsters, including flow cytometric analyses and detection of cytokine production at the protein level. Further investment in the development of these types of reagents is warranted as the hamster model of SARS-CoV-2 infection has shown great utility.

Several promising discoveries have recently been achieved using the widely adopted golden hamster model of SARS-COV-2 infection to examine viral pathogenesis^69,70^, assess immunity^71^, and evaluate of antivirals^72^, vaccines^73^, and therapeutics^74^. Taken together, the data presented here indicate that differences in immune responses between route of exposure, sex and age could significantly affect the outcome of SARS-CoV-2 infection in golden hamsters. It will be important to further understand the effects of each of these variables and how they may influence the outcome of pre-clinical challenge studies. Continued analysis of the immune response to SARS-CoV-2 in animal model systems could further inform future clinical interventions that could modulate the host immune to SARS-CoV-2 infection.

## Figure Legends

**Extended Data Figure 1. Virus distribution in the nasal turbinates**.

In situ hybridization (ISH) using antisense probes that detect the SARS-CoV-2 genome/mRNA on nasal turbinates tissue of uninfected and SARS-CoV-2-infected hamsters (*Mesocricetus auratus*) exposed by the indicated routes of infection at 5 dpi. Six-week-old mixed female and male golden hamsters were inoculated with 10^5^ TCID^50^ of SARS-CoV-2 by a low (20 µl, i.n.L) or high (100 µl, i.n.H) volume intranasal route of administration or an intragastric (i.g.) route of administration and compared to age-matched uninfected controls. Positive detection of viral genomic RNA/mRNA is indicated by magenta staining. The magnification is 10x. Scale bars = 100 µm.

**Extended Data Figure 2. Hematological levels and serum biochemistry in SARS-CoV-2-infected golden Syrian hamsters**.

Six-week-old male and female hamsters (*Mesocricetus auratus*) were inoculated with 10^5^ TCID50 of SARS-CoV-2 by a low volume intranasal (orange diamonds), high volume intranasal (red circles) or intragastric (teal squares) route of administration. The high volume intranasal group data is further broken down by sex (female, lavender; male, dark blue) and compared with an additional group of 20-week-old males exposed to 1×10^5^ TCID^50^ of SARS-CoV-2 a high volume intranasal route of exposure (light blue squares). Hematological levels and serum biochemistry were measured in uninfected and SARS-CoV-2-infected hamsters, including (**a**) white blood cell counts, (**b**) lymphocyte counts, (**c**) neutrophil counts, (**d**) the neutrophil-to-lymphocyte ratio, (**e**) alanine aminotransferase (ALT), (**f**) blood albumin (ALB), and (**g**) blood urea nitrogen (BUN). Bars indicate mean, error bars indicate SEM. a-d, n = 6, 5, 9, 5, 5, 4, and 4 for uninfected, i.n.L, i.n.H, i.g., i.n.H-female, i.n.H-male, and i.n.H-older males, respectively. e-f, n = 4, 5, 10, 5, 5, 5, and 4 for uninfected, i.n.L, i.n.H, i.g., i.n.H-female, i.n.H-male, and i.n.H-older males, respectively. * = *P* < 0.05, ** *P* = 0.01, *** *P* = 0.001, **** = *P* < 0.0001, ns = *P* > 0.05; unpaired student t test (a-g).

**Extended Data Figure 3. Humoral immune response to SARS-CoV-2 infection at 21 dpi**.

Six-week-old male and female golden hamsters (*Mesocricetus auratus*) were inoculated with 10^5^ TCID^50^ of SARS-CoV-2 by a low volume intranasal (orange diamonds), high volume intranasal (red circles) or intragastric (teal squares) route of administration. The high volume intranasal group data is further broken down by sex (female, lavender; male, dark blue) and compared with an additional group of 20-week-old males exposed to 1×10^5^ TCID^50^ of SARS-CoV-2 by a high volume intranasal route of exposure (light blue squares). **a**, IgG antibody response against

SARS-CoV-2 spike antigen were assessed by ELISA using serum collected at 21 dpi. **b**, Neutralizing antibody against SARS-CoV-2 was measured by PRNT^90^ using serum collected at 21 dpi.. * = *P* < 0.05, ** *P* = 0.01, *** *P* = 0.001, **** = *P* < 0.0001, ns = *P* > 0.05; unpaired student t test (a-g).

**Extended Data Figure 4. SARS-CoV-2 re-challenge of i.n.H SARS-CoV-2-infected male golden Syrian hamsters at 133/140 dpi**.

Six-week-old male golden hamsters (*Mesocricetus auratus*) were inoculated with 10^5^ TCID^50^ of SARS-CoV-2 by a high volume intranasal (i.n.H) route of administration. Hamsters were monitored for 133 (n = 5) or 140 (n = 14) days at which point they were re-challenged with the same strain of SARS-CoV-2. **a**, IgG antibody response against SARS-CoV-2 spike antigen were assessed by ELISA using serum collected at 133 or 140 dpi, prior to re-challenge. The hamsters were monitored and weighted daily (**b**) until 5 days after re-infection at which point tissues were obtained. The viral burden in the **c**) nasal turbinates, **d**) proximal lungs **e**) distal lung, and **f**) small intestines were determined by infectious TCID50 assay.

## Methods

### Ethics statement

The experiments described were carried out at the National Microbiology Laboratory (NML) of the Public Health Agency of Canada. All experiments performed under Animal User Document H-20-006, approved by the Animal Care Committee at the Canadian Science Center for Human and Animal Health in accordance with the guidelines provided by the Canadian Council on Animal Care. All procedures were performed under inhalation anesthesia using isoflurane. All efforts were made to minimize animal suffering and to reduce the number of animals used. All infectious work was performed under biosafety level 3 (BSL-3) conditions or higher.

### Cells and Viruses

Vero cells (ATCC) were cultured in Minimal Essential Medium (MEM) (Hyclone) supplemented with 5% Bovine Growth Serum Supplemented Calf (Hyclone) and L-glutamine. Cells were cultured at 37°C with 5% CO^2^. SARS-CoV-2 (Canada/ON-VIDO-01/2020; EPI_ISL_425177) was isolated from a positive patient sample and stocks of virus were grown in VeroE6 cells. Virus stocks were titered by TCID^50^ assay before being used for subsequent *in vivo* experiments. All virus used for *in vivo* experiments was from passage 2.

### Hamster challenge and re-challenge experiments

Groups of male or female golden Syrian hamsters were anaesthetized and exposed to 1×10^5^ TCID^50^ SARS-CoV-2 by an intranasal route of inoculation with the inoculum distributed evenly into both nares in a volume of 20 µl (intranasal low volume) or in a volume of 100 µl to allow for inhalation of the inoculum (intranasal high volume), or by an intragastric route of infection via oral gavage. Mock infected animals were inoculated by an intranasal route with with 100 µL of neat DMEM. Animals were monitored daily for clinical signs of disease including lethargy, hunched posture, inactivity, and labored breathing. Temperatures were measured daily via implanted transponders, and all animals were weighed once daily until 21 dpi and again on 28 dpi. Animals meeting the pre-determined endpoint criteria were anaesthetized and humanely euthanized. Previously exposed hamsters were re-infected with the same strain of SARS-CoV-2 at 81, 133, or 140 dpi and were necropsied at 5 dpi to assess the viral burden in the tissues, viral shedding, and the humoral immune response.

### Hematology and Blood biochemistry

Complete blood counts (CBC) and hematological analysis was performed using a VetScan HM5 (Abaxis Veterinary Diagnostics), as per manufacturer’s instructions. Serum biochemistry values were determined with a VetScan VS2 (Abaxis Veterinary Diagnostics) using complete diagnostic profile disks according to manufacturer’s instructions. Blood and serum obtained from uninfected hamsters was used to establish baseline values.

### Measurement of viral burden in the tissues

For measurement of viral titers in the blood and tissues of infected animals, TCID^50^ assays were performed. Following necropsy, blood and tissue samples were frozen at -80°C for storage. For infectious assays, tissue samples were thawed and placed in MEM, supplemented with 1x L-glutamine and 1% FBS, and homogenized with 5 mm stainless steel beads in a Bead Ruptor Elite Tissue Homogenizer (Omni). Homogenates were clarified by centrifugation at 1500 x g for 10 minutes and ten-fold serial dilutions of tissue homogenates were made in MEM. GI tract tissues were additionally processed to minimize the presence of fecal bacteria: feces were carefully removed from the GI tract lumen and dilutions and infectious assays were carried out in MEM supplemented with 1x L-glutamine, 1% FBS, and a 2x dose of penicillin/streptomycin. Dilutions were added to 90-100% confluent Vero cells in triplicate wells and cytopathic effect was read at 5 dpi. TCID^50^ values per mL or gram of tissue were calculated using a modified Reed and Muench method^75^.

For determination of viral RNA, collected tissues were stored in RNAlater. RNA was extracted using an RNeasy mini plus kit (Qiagen), according to manufacturer’s instructions. For viral RNA present in blood, RNA was extracted using a viral RNA mini kit (Qiagen). RT-qPCR detection of SARS-CoV-2 was performed on a QuantStudio 5 instrument (Applied Biosystems) using a TaqPath 1-step RT-qPCR Master Mix (Applied Biosystems) and primers specific for the E gene of SARS-CoV-2 as per the diagnostic protocol recommended by the World Health Organization (Forward – ACAGGTACGTTAATAGTTAATAGCGT; Reverse – ATATTGCAGCAGTACGCACACA; Probe – FAM-ACACTAGCCATCCTTACTGCGCTTCGBBQ). Oligonucleotide concentrations were 400nM for the primers and 200nM for the probe. RT-qPCR stages were as follows: UNG incubation (25°C for 2 minutes), reverse transcription (53°C for 10 minutes), polymerase activation (95°C for 2 minutes), followed by amplification (40 cycles of 95°C for 3 seconds and 60 °C for 30 seconds).

### Detection of virus in mucosal swab samples

Oropharyngeal and rectal swabs as well as nasal washes were obtained from animals during necropsy. Swabs were stored in MEM + 2% penicillin-streptomycin. Prior to titration procedures, tubes containing swabs were vortexed and centrifuged briefly. For viral RNA detection, 140µL of the medium containing the swab was used for viral lysis and extraction using a viral RNA mini kit as above.

### Transcriptional profiling of host responses

Tissue RNA was extracted as described above using an RNeasy mini plus kit, which includes a genomic DNA eliminator step. Host mRNA expression of various genes including IL1β, IL6, TNFα, IL2, IFNγ, IL4, IL10, FoxP3, STAT2, and Mx2 was quantified as described previously using RPL18 as an internal reference gene^76^. RT-qPCR was performed using a TaqPath 1-step RT-qPCR kit as described above.

### SARS-CoV-2-S-specific enzyme-linked immunosorbent assay (ELISA)

Ninety-six-well flat-bottom high-binding microplates (Corning, New York, USA) were coated with recombinant SARS-CoV-2 spike protein in PBS at concentration of 25 ng/well overnight at 4°C. The next day, plates were washed four times with PBS-T (PBS + 0.1% Tween 20) and then blocked with blocking buffer (PBS-T + 5% skim milk powder) for 1 hour at 37°C. Following blocking, plates were washed four times with PBS-T and serum samples from infected or mock infected hamsters were serially diluted and added to the plates in triplicate for 1 hours at 37°C. Plates were then washed with PBS-T four times and secondary peroxidase AffiniPure Goat Anti-Syrian Hamster IgG (H+L) (Jackson IR, #107-035-142) was added to the plates at a dilution of 1:1000 for 1 hour at 37°C. Plates were once again washed with PBS-T and ABTS substrate was added to the plates for 30 minutes at room temperature and then OD^405nm^ readings were taken.

### Plaque reduction virus neutralization test (PRNT-90)

Hamster serum samples were collected and stored at -80°C. SARS-CoV-2 stocks were titrated and used in the PRNT-90^77^. In short, serum was heat-inactivated at 56°C for 30 minutes and diluted 2-fold (dilution range 1:40 to 1:1280) in DMEM supplemented with 2% FBS. Diluted sera was incubated with 50 plaque forming units of SARS-CoV-2 at 37°C with 5% CO_2_ for 1 hour. The sera-virus mixtures were added to VeroE6 cells at 100% confluence in 24-well plate format, followed by incubation at 37 °C and 5% CO_2_ for 1 hour. After adsorption, 1.5% carboxymethylcellulose diluted in MEM supplemented with 4% FBS, L-glutamine, non-essential amino acids, and sodium bicarbonate was added to each well. Plates were then incubated at 37°C and 5% CO_2_ for 72 hours. The liquid overlay was removed and the cells were fixed with 10% neutral-buffered formalin for 1 hour at room temperature. The monolayers were then stained with 0.5% crystal violet for 10 minutes and washed with 20% ethanol. Plaques were counted and compared to a 90% neutralization control. The PRNT-90 endpoint titre was defined as the highest dilution of serum resulting in a 90% reduction of plaques. PRNT-90 titers ≥1:40 were considered positive for neutralizing antibodies.

### Histopathology and vRNA in situ hybridization

Tissues were fixed in 10% neutral phosphate buffered formalin for a minimum of 7 days. Routine processing was carried out and tissue samples were sectioned at 5 µm. A set of slides was stained with hematoxylin and eosin for histopathologic examination. RNA in situ hybridization (ISH) was carried out using RNAscope 2.5 HD Detection Reagent-Red (Advanced Cell Diagnostics), according to the manufacturer’s instructions. Briefly, formalin-fixed, paraffin-embedded tissue samples were mounted on slides, baked in a dry oven for 1 hour at 60°C, and deparaffinized. Tissue sections were then pre-treated with RNAscope H^2^O^2^ for 10 minutes at room temperature, and target retrieval was carried out using the RNAscope Target Retrieval Reagent for 15 minutes. RNAscope Protease Plus Reagent was then applied for 15 minutes at 40°C. The probes targeting SARS-CoV-2 RNA (V-nCoV2019-S probe, ref#848561) or anti-genomic RNA (V-nCoV2019-S-sense ref#845701) were designed and manufactured by Advanced Cell Diagnostics, and the negative probe was also obtained from Advanced Cell Diagnostics (Reference # 310034). The stained tissues were counterstained with Gills I Hematoxylin. The final images were captured using a light microscope equipped with a digital camera.

### Data analysis

Results were analyzed and graphed using Prism 8 software (Graphpad Software). As appropriate, statistical analyses were performed using ANOVA with multiple comparison correction, the multiple t test, or the unpaired t test with Welch’s correction or Mann-Whitney test.

### Data availability

All relevant data are available from the authors upon request.

