## Supplementary Figure 1 for "Exposure route, sex, and age influence disease outcome in a golden Syrian hamster model of SARS-CoV-2 infection"

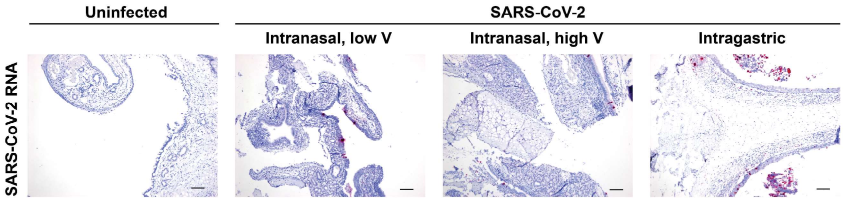


**Extended Data Figure 1. Virus distribution in the nasal turbinates**. In situ hybridization (ISH) using antisense probes that detect the SARS-CoV-2 genome/mRNA on nasal turbinates tissue of uninfected and SARS-CoV-2-infected hamsters exposed by the indicated routes of infection at 5 dpi. Six-week old mixed female and male golden hamsters were inoculated with $10^5$ $TCID_{50}$ of SARS-CoV-2 by a low (20 µl, i.n.L) or high (100 µl, i.n.H) volume intranasal route of administration or an intragastric (i.g.) route of administration and compared to age-matched uninfected controls. Positive detection of viral genomic RNA/mRNA is indicated by magenta staining. The magnification is 10x. Scale bars = 100 µm.
