## Supplementary Figure 2 for "Exposure route, sex, and age influence disease outcome in a golden Syrian hamster model of SARS-CoV-2 infection"

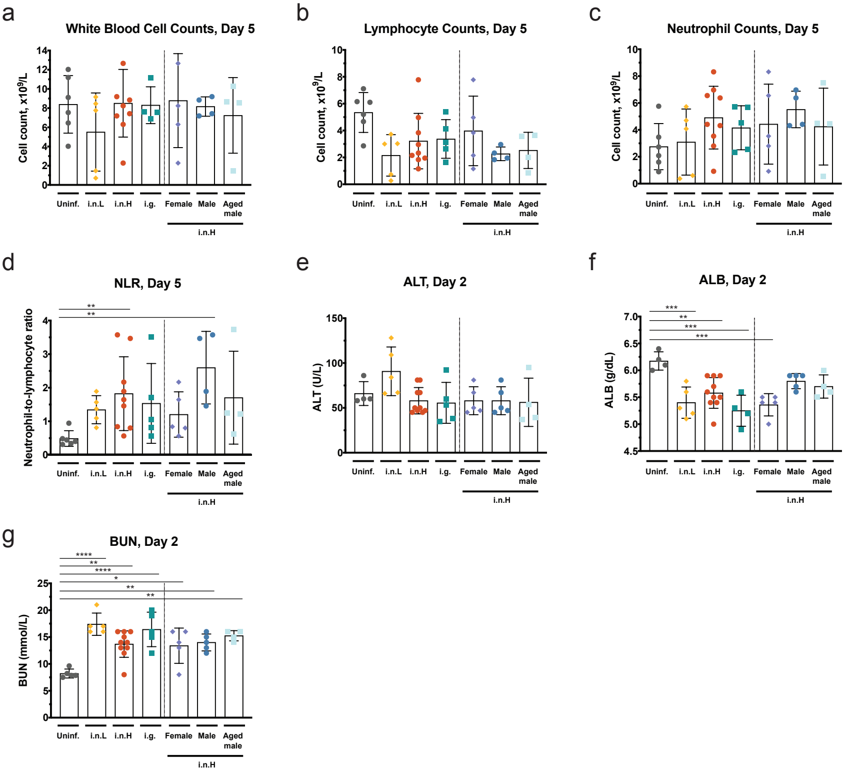


**Extended Data Figure 2. Hematological levels and serum biochemistry in SARS-CoV-2-infected golden Syrian hamsters.** Six-week-old male and female hamsters were inoculated with $10^5$ $TCID_{50}$ of SARS-CoV-2 by a low volume intranasal (orange diamonds), high volume intranasal (red circles) or intragastric (teal squares) route of administration. The high volume intranasal group data is further broken down by sex (female, lavender; male, dark blue) and compared with an additional group of 20-week-old males exposed to the same dose by a high volume intranasal route of exposure (light blue squares). Hematological levels and serum biochemistry were measured in uninfected and SARS-CoV-2-infected hamsters, including (a) white blood cell counts, (b) lymphocyte counts, (c) neutrophil counts, (d) the neutrophil-to-lymphocyte ratio, (e) alanine aminotransferase (ALT), (f) blood albumin (ALB), and (g) blood urea nitrogen (BUN). Bars indicate mean, error bars indicate SEM. a-d, n = 6, 5, 10, 5, 5, 4, and 4 for uninfected, i.n.L, i.n.H, i.g., i.n.H-female, i.n.H-male, and i.n.H-older males, respectively. e-f, n = 4, 5, 10, 5, 5, 5, and 4 for uninfected, i.n.L, i.n.H, i.g., i.n.H-female, i.n.H-male, and i.n.H-older males, respectively. * = $P < 0.05$, ** $P = 0.01$, *** $P = 0.001$, **** = $P < 0.0001$, ns = $P > 0.05$; unpaired student t test (a-g).
