## Supplementary Figure 3 for "Exposure route, sex, and age influence disease outcome in a golden Syrian hamster model of SARS-CoV-2 infection"

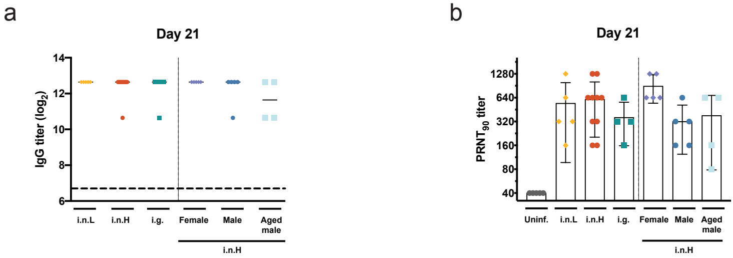


**Extended Data Figure 3. Humoral immune response to SARS-CoV-2 infection at 21 dpi.** Six-week-old male and female golden hamsters were inoculated with $10^5$ $TCID_{50}$ of SARS-CoV-2 by a low volume intranasal (orange diamonds), high volume intranasal (red circles) or intragastric (teal squares) route of administration. The high volume intranasal group data is further broken down by sex (female, lavender; male, dark blue) and compared with an additional group of 20-week-old males exposed to the same dose by a high volume intranasal route of exposure (light blue squares). **a**, IgG antibody response against SARS-CoV-2 spike antigen were assessed by ELISA using serum collected at 21 dpi. **b**, Neutralizing antibody against SARS-CoV-2 was measured by $PRNT_{90}$ using serum collected at 21 dpi.
