## Supplementary Figure 4 for "Exposure route, sex, and age influence disease outcome in a golden Syrian hamster model of SARS-CoV-2 infection"

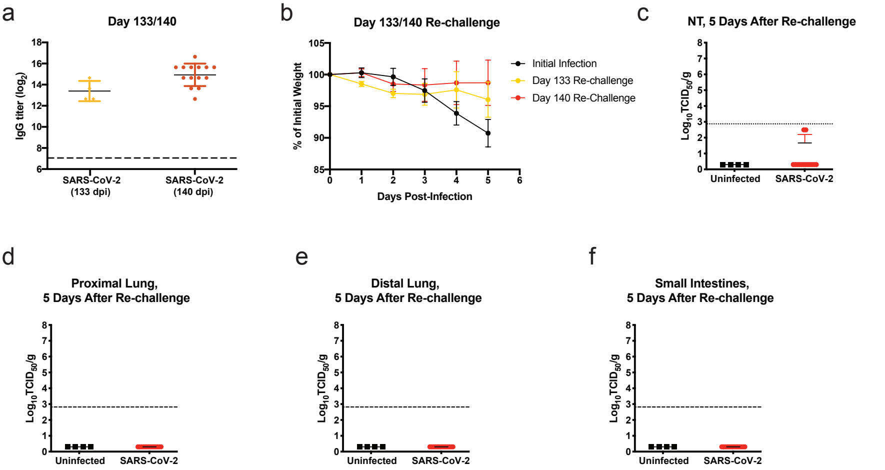


**Extended Data Figure 4. SARS-CoV-2 re-challenge of i.n.H SARS-CoV-2-infected male golden Syrian hamsters at 133/140 dpi.** Six-week-old male golden hamsters were inoculated with $10^5$ $TCID_{50}$ of SARS-CoV-2 by a high volume intranasal (i.n.H) route of administration. Hamsters were monitored for 133 or 140 days at which point they were re-challenged with the same strain of SARS-CoV-2. **a**, IgG antibody response against SARS-CoV-2 spike antigen were assessed by ELISA using serum collected at 133 or 140 dpi, prior to re-challenge. The hamsters were monitored and weighted daily (**b**) until 5 days after re-infection at which point tissues were obtained. The viral burden in the **c**) nasal turbinates, **d**) proximal lungs **e**) distal lung, and **f**) small intestines were determined by infectious $TCID_{50}$ assay.
